# Atomistic Simulation of Blood–Brain Barrier Permeability of Propolis-Derived Natural Compounds

**DOI:** 10.64898/2026.06.08.730943

**Authors:** Vipul Kumar, Sunil C. Kaul, Renu Wadhwa, Durai Sundar

**Author notes:** Department of Bioengineering and Therapeutic Sciences, University of California, San Francisco, San Francisco, California, USA.

## Abstract

The ability of small molecules to cross the blood–brain barrier (BBB) remains a major bottleneck in neurotherapeutic development. While experimental assays and machine learning approaches provide approximate permeability estimates, they lack atomistic insight into the underlying transport mechanisms. Here, we employ all-atom molecular dynamics simulations of a compositionally realistic BBB lipid bilayer to characterize the passive permeation of two bioactive propolis-derived compounds, Caffeic Acid Phenethyl Ester (CAPE) and Artepillin-C (ARC). Using steered molecular dynamics and umbrella sampling, we computed free energy profiles, diffusion coefficients, and permeability metrics across the membrane. CAPE encounters a modest barrier at the lipid headgroup region but minimal resistance within the hydrophobic core, resulting in a low free energy barrier (∼2–3 kcal/mol) and favorable permeability (logP_eff ≈ 0.28). In contrast, ARC exhibits a substantial energetic barrier within the membrane core, leading to high resistivity and strongly unfavorable permeability (logP_eff ≈ −10.91). The heterogeneous lipid model reproduces experimentally consistent membrane properties and reveals how lipid composition modulates transport energetics. These findings provide mechanistic insight into BBB permeability and demonstrate the utility of atomistic simulations for guiding the design of neuroactive therapeutics.

## 1. Introduction

The ability of drug molecules to cross biological membranes is a fundamental requirement for achieving therapeutic efficacy. In the context of central nervous system (CNS) disorders, this challenge is amplified by the presence of the blood–brain barrier (BBB), a highly selective interface that tightly regulates molecular transport into the brain (Sun et al. 2022). Despite promising target engagement in early-stage studies, many drug candidates fail in later stages due to insufficient membrane permeability and poor bioavailability, making permeability a critical determinant of drug success. Molecular transport across membranes occurs primarily via two mechanisms: active transport, which requires energy input, and passive diffusion, driven by concentration gradients. For most small, drug-like molecules, passive diffusion is the dominant pathway. Consequently, understanding the physicochemical and structural factors governing passive permeation is essential for rational drug design (Sharifian Gh 2021).

A range of experimental and computational approaches have been developed to estimate BBB permeability. Experimental techniques such as parallel artificial membrane permeability assays (PAMPA) and immobilized artificial membrane chromatography provide useful approximations but are often labor-intensive and limited in mechanistic resolution (Ottaviani et al. 2006) (Ong et al. 1996). Computational approaches, including quantitative structure– activity relationship (QSAR) models and machine learning methods, offer rapid predictions based on molecular descriptors; however, they lack direct insight into the molecular interactions governing transport (Toropov et al. 2017; Faramarzi et al. 2022; Zhang et al. 2008). In contrast, all-atom molecular dynamics (MD) simulations enable explicit modeling of membrane environments and provide a physically grounded framework for calculating free energy profiles, diffusion behavior, and permeability at atomic resolution (Carpenter et al. 2014) (Radhakrishnan et al. 2022; Wadhwa et al. 2021). Although it is a computationally expensive process, it allows us to calculate the diffusion rate at the atomic level. While many MD studies of membrane permeability employ simplified lipid bilayers composed predominantly of phosphatidylcholine. While such models are useful for comparative analysis, they do not adequately capture the compositional complexity of biological membranes. The BBB, in particular, is characterized by a distinct lipid composition, including elevated levels of sphingomyelin and cholesterol and reduced phosphatidylcholine content, which significantly influence membrane packing, fluidity, and transport energetics. Therefore, incorporating realistic lipid composition is essential for accurately modeling permeation processes across the BBB (Carpenter et al. 2014; Boggara and Krishnamoorti 2010; Orsi et al. 2009). These MD simulation studies could not completely mimic the actual environment, however relevant to calculate the small molecules’ relative permeability.

In this study, we have tried to make a complex bilayer lipid membrane instead of taking homogeneous lipids to study the permeation of the natural compounds through the BBB. The major challenge in neurotherapeutics is delivering the drugs inside the brain tissue. The drug candidate which intends to interact with the target nervous system should pass through a very complex barrier, generally known as BBB. The BBB is made from endothelial cells that make the wall of cerebral capillaries, which serves as both physical and biological barriers to control the brain environment (Daneman and Prat 2015). The major difference between normal endothelial cells and brain capillaries endothelial cells is the composition of lipids in the plasma membrane and tight junctions between the endothelial cells (Stamatovic et al. 2008). Claudin-5 membrane proteins constitute the tight junctions (TJ) that act as gatekeepers of molecular transport in the BBB. Apart from these differences, two other types of cells, pericytes and astrocytes, work as a barrier and cover almost 80% of the endothelial cells (Campbell et al. 2014). The lipid composition of brain micro-vessel endothelial cells is the primary determinant of the BBB. The amount of sphingomyelin is approximately 3-fold higher, and phosphatidylcholine content is decreased by approximately one-third, compared with other endothelial cells (Tewes and Galla 2001). Therefore the BBB model built for this study is similar to the experimentally known lipid profiles (detailed in the method section).

In this study, we tested the permeability of 5 molecules (3 controls) and (2 test molecules) (**Figure 1 A-E**). The two test molecules chosen for the permeability prediction are well-known active compounds from honeybee propolis: Caffeic Acid Phenethyl Ester (CAPE) and Artepillin-C (ARC). These natural products are known the neuroprotective activities and the preclinical evidence of the molecules targeting the CNS targets is also available in the literature (Kaul et al. 2021; Kulkarni et al. 2021). However, to the best of our knowledge, the permeability of these compounds through BBB is not studied yet.

**Figure 1.**
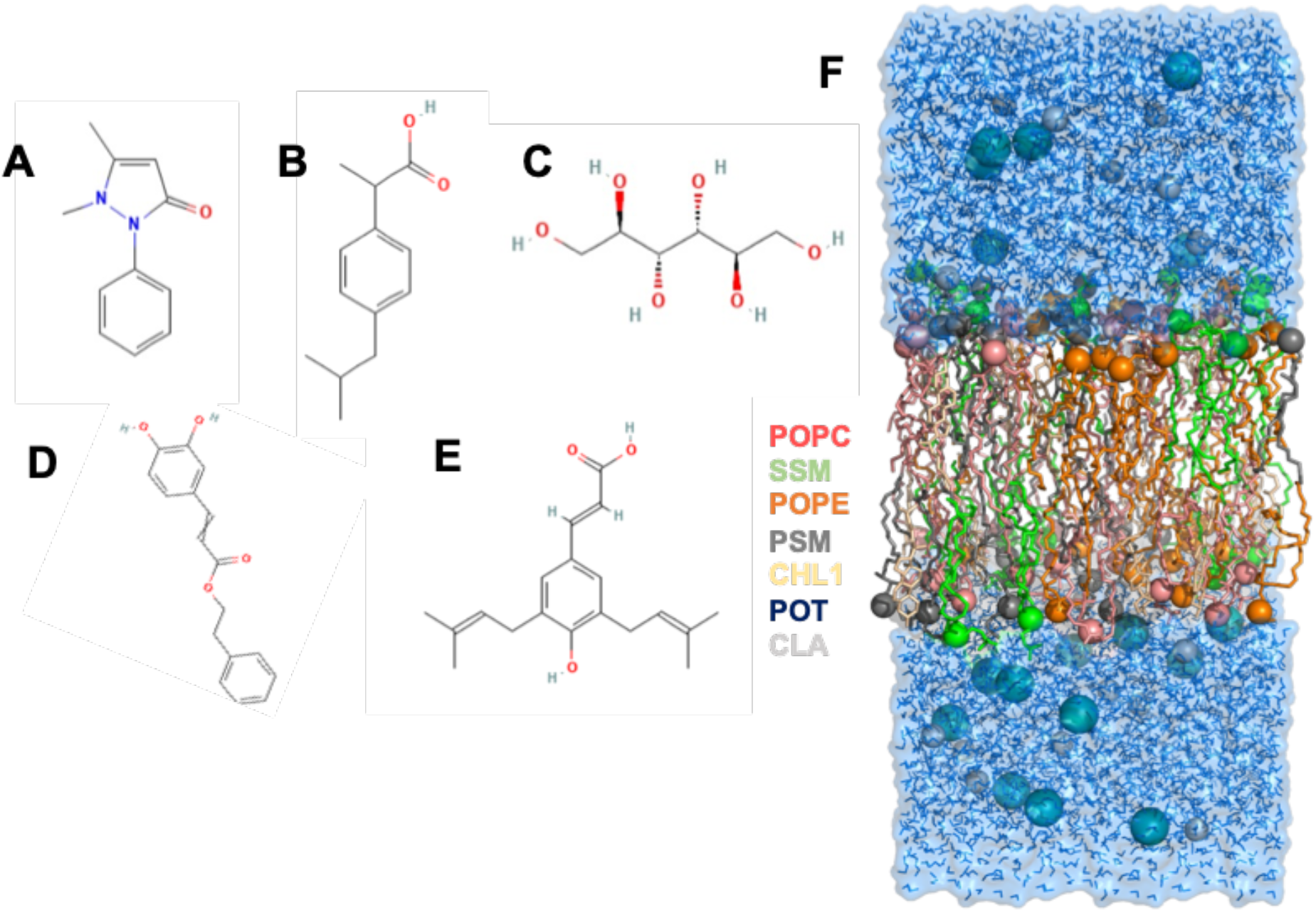
The 2D images of the small molecules studied and the 3D representation of the modelled BBB bilayer model. (A) Antipyrine (B) Ibuprofen (C) Mannitol (D) CAPE (E) ARC (F) The equilibrated BBB model contains POPC in red, POPE in orange, PSM in grey, cholesterol in yellow, and SSM in green.

We have computed these compounds’ diffusion rate and logP value using the steered molecular dynamics simulation and umbrella sampling technique. We have compared the computed logP with the experimental logBBB. Logp≥0 has been taken as BBB^+^ (Permeable) and logp<0 as BBB^-^ (Non-Permeable).

## 2. Material and Methods

### 2.1 Modelling of BBB lipid bilayer

The BBB model was built similarly to the experimentally known lipid profiles using CHARMMGUI (Jo et al. 2008). In the rectangular system, the thickness was 40 Å (minimum water height on top and bottom of the system), and the hydration number chosen was 50 (number of water molecules per lipid molecule). In this model, 50 lipid molecules at each leaflet were added to make it symmetric. Lipid molecules constituted of 10 Cholesterol molecules, 13 phosphatidylcholines (POPC) molecules, 10 Phosphatidylethanolamine (POPE), and 17 sphingomyelins (PSM and SSM) molecules were added at each leaflet (**Table1**). The total area of each leaflet was 2816.8 Å^2^. Then K^+^ and Cl^-^ were added to neutralize the system (0.15 M). The complete system contained around ∼ 32168 atoms, 50 lipid molecules at each leaflet and 6748 water molecules (TIP3P). The dimensions of the system were 53.07*53.07*120 Å^3^ (Lee et al. 2016).

**Table1.**
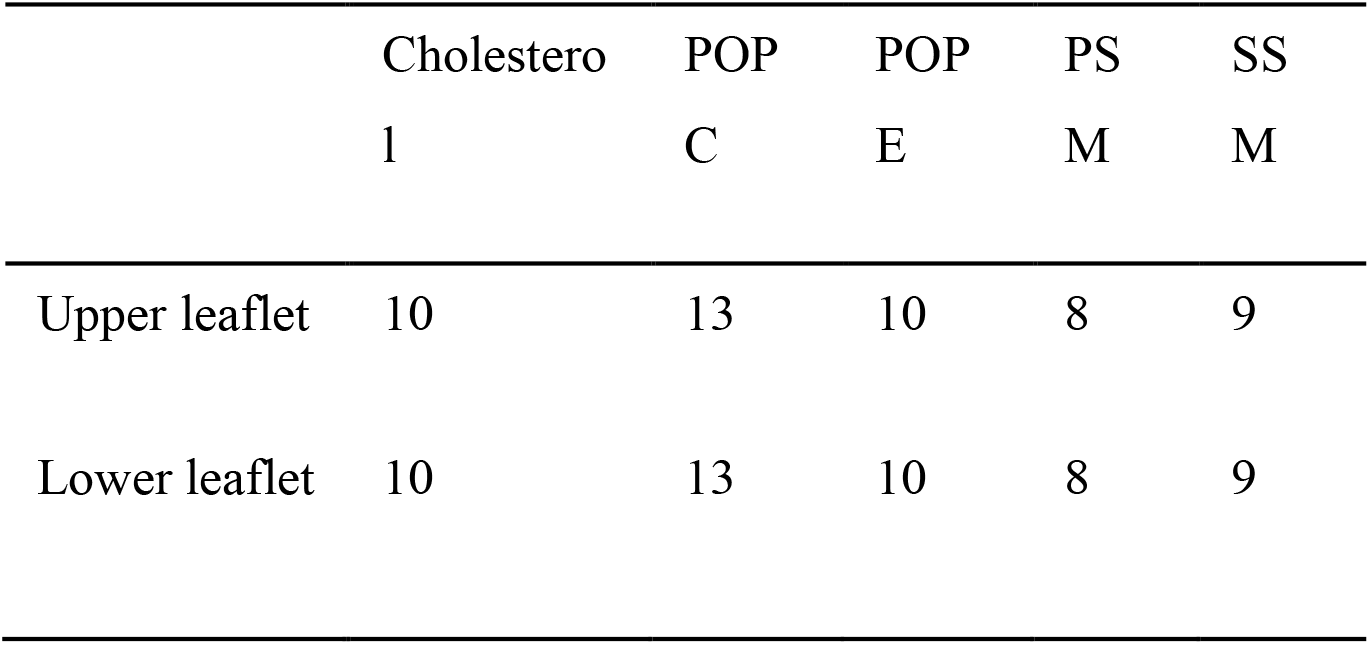
The distribution of different lipid molecules in the BBB model.

### 2.2 Minimization, equilibration and Production MD of the lipid bilayer system

The energy minimization, equilibration and production run of the built BBB model system were done using GROMACS 2020 (Abraham et al. 2015). The detailed protocol of these methods have been reported elsewhere (Radhakrishnan et al. 2022). The production simulation was used for calculating all the properties of the bilayer membrane system (Smith et al. 2019).

### 2.3 Properties of the lipid bilayer

The MEMBPLUGIN in VMD was used to calculate the various properties of the modelled lipid bilayer to validate the modelled membrane (Guixà-González et al. 2014). Firstly, the density profile of all the lipids, water and the whole system was calculated through gmx_density using GROMACS2020 (Abraham et al. 2015). Then the area per lipid and order parameter of the lipid tails were calculated.

### 2.4 Free energy calculations

To calculate the PMF (potential of Mean Force), the steered molecular dynamics simulations were first performed to sample the energy and configuration of drug molecules along the bilayer. As the bilayer is identical (i.e., the lower leaflet matches the upper leaflet), the calculations have only been performed on molecules passing through one side of the membrane (Park and Schulten 2004). The forcefield parameter for the small molecules in the study was generated through CgenFF (Vanommeslaeghe et al. 2010). The drug molecules were inserted in the water ∼ 2-3Å above the lipid head group. All the steered MD simulations were done in triplicates, so the molecules were inserted randomly in the X-Y plane while keeping the Z dimension the same (height from the lipid molecule) to minimize the interaction bias between the same lipid and drug molecules. The average values have been presented in the results. Following the insertion of the drug molecules, the systems were minimized for 5000 steps using the steepest descent algorithm. The minimization was followed by 2ns of equilibration in the NPT ensemble while putting the position constraints (1000 kJ/mol/nm^2^) on the drug molecules in the Z-direction. The steered MD simulations were performed in the NPT ensemble using GROMACS pulling code. The Noose-Hoover with a time constant of 1 ps was used to maintain the 303 K temperature, and the Parrinello-Rahman barostat with a time constant of 5ps was used to maintain the 1 atm pressure during the simulation. The drug compounds were pulled towards the lipid bilayer’s hydrophobic core (centre) at 0.001 nm/ps speed using the force constant of 1000 kJ/mol/nm^2^ (Abraham et al. 2015).

Further, the coordinates were sampled for every 0.2 nm, dividing the whole coordinates into 25 windows through umbrella sampling (US) (Torrie and Valleau 1977). Then each window was simulated for 30ns in an NPT ensemble (303 K and 1 atm) with a force constant of 500 kJ/mol/nm^2^. The first 10ns of each window was regarded as the equilibration time given to the compound to orient correctly in the system. The last 20ns of each window are used for further analysis. Temperature and pressure were maintained by Noose-Hoover and Parrinello-Rahman algorithms, respectively. The weighted histogram analysis method (WHAM) within the GROMACS was used to calculate each molecule’s PMF free energy profile based on the US data (Abraham et al. 2015; Kumar et al. 1992).

### 2.5 The diffusion and the permeability of the small molecules

The permeability of the small molecules from the lipid bilayer can be computed by using the information of the PMF and diffusion coefficient profile of the molecules throughout the reaction coordinates.

The position-specific diffusion coefficients were calculated using umbrella sampling data by GROMACS gmx_msd code (Abraham et al. 2015). This method provides an easy way to compute the diffusion coefficient using the Einstein relation (Allen and Tildesley 1987). Further, the resistivity of the small molecules was calculated through PMF and diffusion profile through the following equations (Marrink and Berendsen 1994):

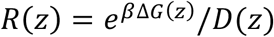

Here, R is the resistivity along the Z axis, *β is the B*oltzmann constant, which is equal to 1/*K*_’_*T*, D is the diffusion coefficient.

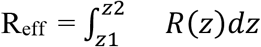

R_eff_ is effective resistivity.

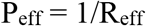

P_eff_ is the effective permeability of the compound.

## 3. Results and Discussion

The lipid composition in the human cell membrane is usually a complex mixture of lipids and proteins. For this study, we tried to mimic the lipid environment of the human BBB lipid profile. It is reported that human brain endothelial cell contains 28.7% of POPC, 22.8% of POPE, 33.7% of sphingomyelin and ∼20% of cholesterol. Using this information, we prepared the lipid bilayer model total of 100 lipids, 50 at each leaflet, in the same proportion as reported earlier (Tewes and Galla 2001; Campbell et al. 2014). It contained 10 Cholesterol molecules, 13 POPC molecules, 10 POPE, and 17 sphingomyelins (PSM and SSM) molecules. The prepared model was then minimized and simulated for 500ns (equilibration + production run) before inserting the drug molecules (**Figure 1F**). The fully equilibrated system was then used to calculate membrane properties and compare them to experimental data.

### 3.1 The prepared BBB model properties were consistent with experimental data

The model lipid system should have the desired fluidity. The density profile of each lipid was assessed to verify the localization of lipids in the system. The density of each lipid across the bilayer is shown in **Figure 2 A**. On the X axis, 0 indicates the centre of the hydrophobic core, while a negative value indicates the outer leaflet and a positive value towards the inner leaflet. The peaks at -2 and +2 in the total lipid profile show the polar head group of the lipid molecules. At the same time, the water density is prominent in the periphery of the lipid bilayer. The density profiles were similar to already published data, signifying the correctly equilibrated system for the permeation studies (Wadhwa et al. 2021; Shahane et al. 2019). The membrane thickness was 47.56 ± 0.57 Å, close enough to the previous membrane models (Carpenter et al. 2014; Shahane et al. 2019). Next, the area per lipid was calculated and analyzed. The area per lipid profile (∼60 Å^2^ for POPC and POPE; ∼ 30 Å^2^ for Sphingomyelins and Cholesterol) shown in **Figure 2B**, calculated throughout the simulations, is consistent with the experimentally known data (Mukhopadhyay et al. 2004; Kinnun et al. 2015; Petrache et al. 2000). After area per lipid calculations, the Scd order parameters of the lipids were calculated.

**Figure 2.**
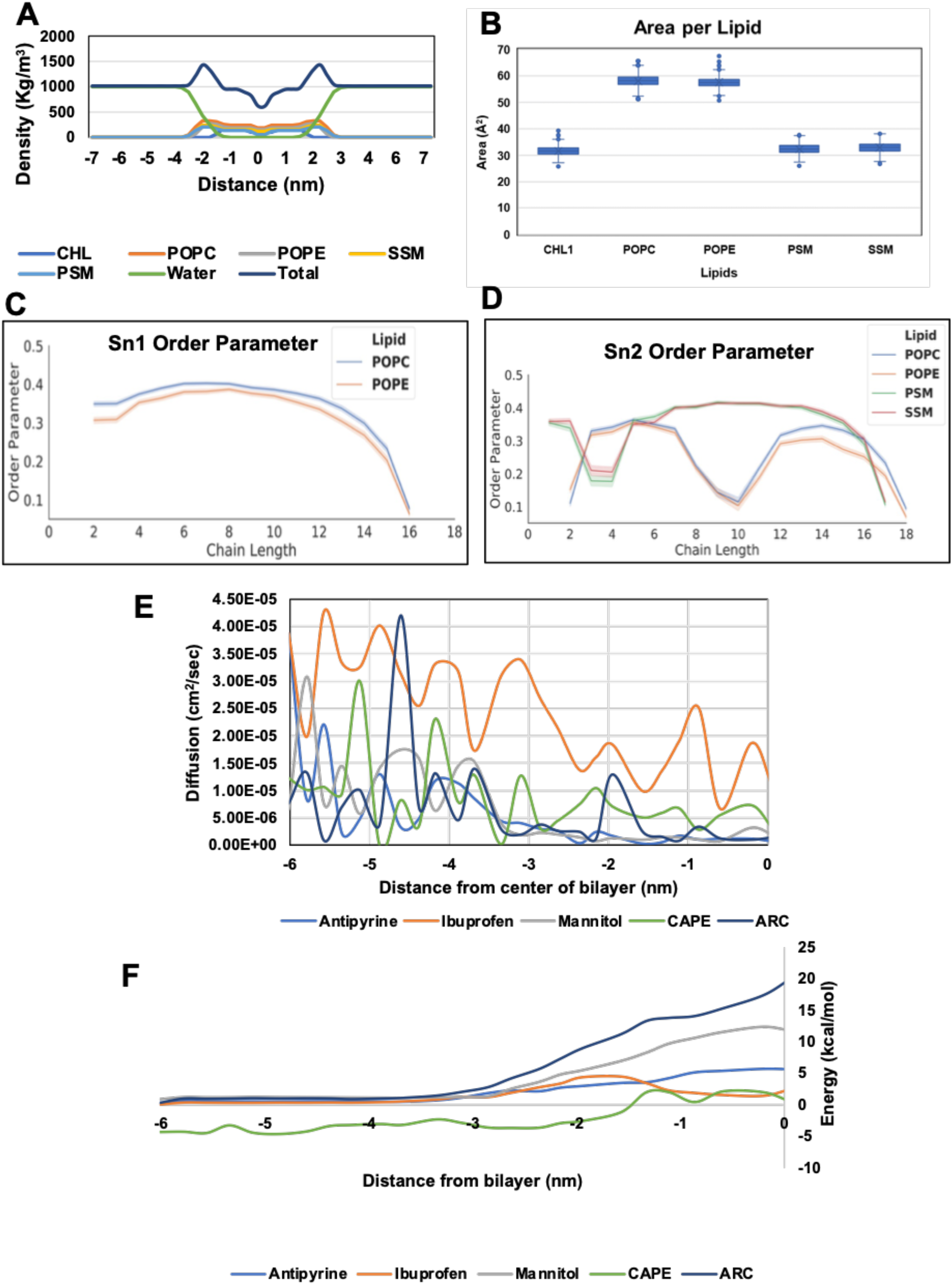
The structural properties of lipid molecules and Diffussion profile of small molecules. (A) The density profile of all the molecules within the system shows the distribution of their localization (0 represents the centre of the bilayer). (B) The average area per lipid profile of the lipid molecules. (C, D) The Sn1 and Sn2 order parameters of the lipid molecules in the BBB model. (E) The diffusion profile shows the higher diffusion of smaller molecules and a higher diffusion rate in the water. (F) The PMF profile shows the highest energy barrier of ARC than CAPE to pass the BBB.

The order parameter of the lipid tails is a measure of the degree of order or alignment of the lipid tails within a membrane. Sn1 and Sn2 tail orders were calculated for POPC and POPE (containing two hydrocarbon chains), while Sn2 tail orders for SSM and PSM. The low order parameter in Sn2 chain C3-4 and C9-10 may be due to the double bond (**Figure 2C-D**). The calculated order parameter data aligned with previously published data (Hofsäss et al. 2003).

### 3.2 Diffusion, PMF and Resistivity Profiles of the small molecules

Steered Molecular Dynamics (SMD) and umbrella sampling were performed to generate the diffusion and PMF profile of the tested small molecules, namely, Antipyrine, Mannitol, Ibuprofen, ARC and CAPE. Firstly diffusion coefficient was calculated for all these compounds, which shows the diffusion rate of these compounds. Smaller molecules tend to have higher diffusion than bigger molecules. Further, it was observed that all the tested molecules had a higher diffusion rate in water than the lipids (**Figure 2E**). The bigger molecules had the lowest diffusion throughout the simulation. Then the PMF was calculated, which indicates the free energy change with respect to the position of the molecules. The highest point in PMF (del G_max_) indicates the highest barrier for a molecule to pass for passage through the membrane. **Figure 2 F** shows the PMF profile of all the tested molecules. Through PMF calculation, it was observed that CAPE had the lowest energy barrier of ∼ 2-3 kcal/mol. Further the CAPE and Ibuprofen showed positive energy (resistance) only when they interacted polar head group (around -2 Å from the centre), while their energy was almost 0 towards the hydrophobic core.

In contrast, the other compounds had the highest barrier towards the hydrophobic core of the lipid bilayer. To show resistance clearly, the resistivity of the compounds was calculated and assessed. The resistivity profile showed that Antipyrine had the highest resistance in the permeation, followed Mannitol and ARC, and they all had the highest resistance towards the hydrophobic core. While Ibuprofen had the highest resistance at the polar head group of the lipids, CAPE had the resistance at the polar head region and middle tail region of the lipid molecules. Also, CAPE was the molecule which had the lowest resistance in the passage among all the tested molecules (**Figure 3A-G**).

**Figure 3.**
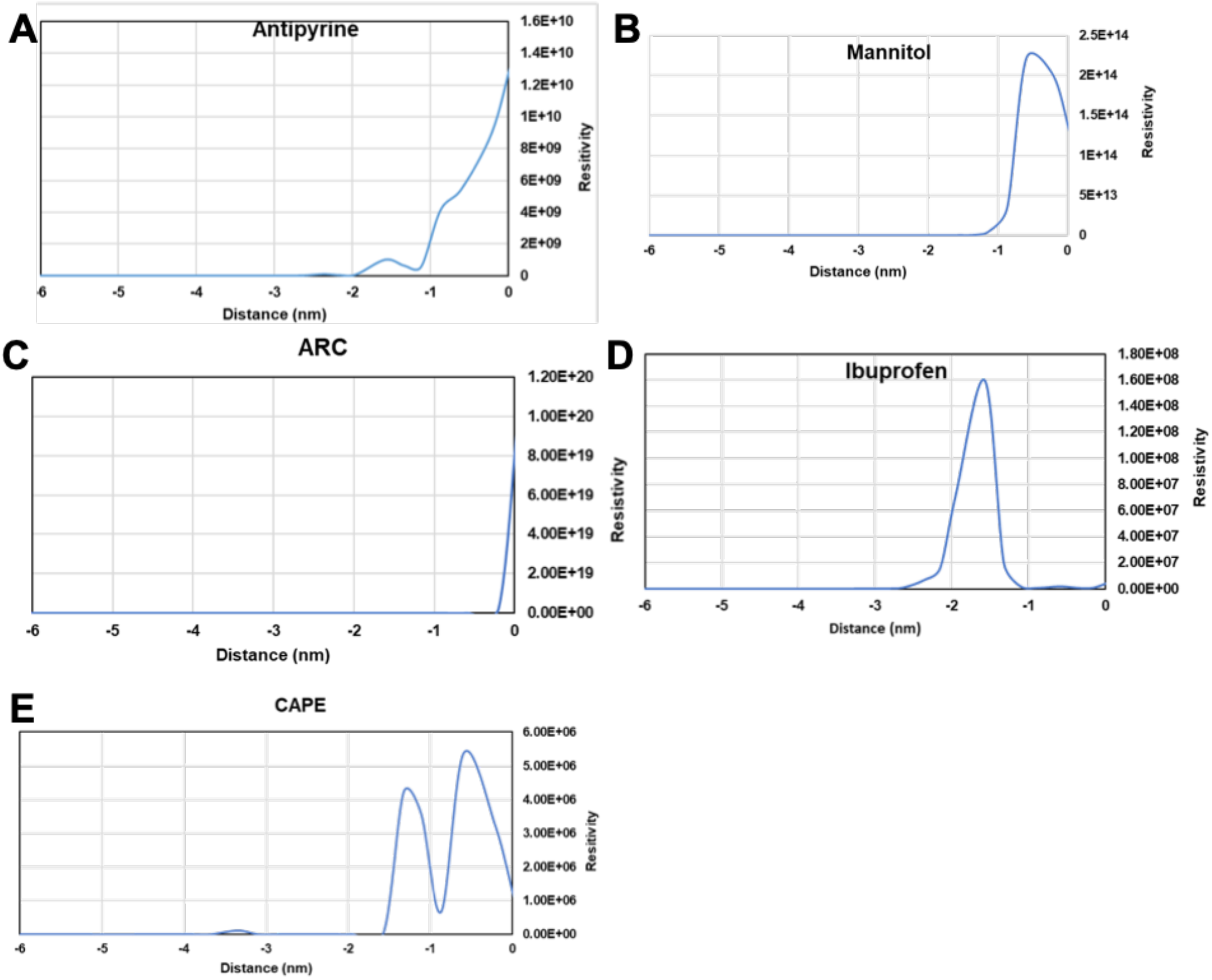
The resistivity profile of the small molecules. (A) Antipyrine (B) Mannitol (C) ARC (D) Ibuprofen (E) CAPE.

### 3.3 CAPE was found to be the permeable passively but not ARC

Through the resistivity calculation, the effective permeability (P_eff_) and log P were calculated, indicating the permeability of the compounds. The log P calculation of Antipyrine, Mannitol and Ibuprofen showed that it was close to the previously reported log P values for BBB and validated the simulation and model (**Table 2**) (Carpenter et al. 2014; Luco 1999). After that, log P of ARC was calculated, which showed the value to be highly negative, -10.91. The log P predicted value indicated that ARC would not permeate the BBB membrane through passive diffusion. On the other hand, the calculated logP value of the CAPE was found to be 0.28, indicating the easy permeation. The Log P and BBB permeability prediction have been shown in **Table 2**. In addition to the energetic differences, the observed permeability trends between CAPE and ARC can be partly explained by their underlying chemical properties. CAPE contains an aromatic ester scaffold that provides moderate lipophilicity, allowing it to interact favorably with both the polar head groups and the hydrophobic core of the membrane. This balance likely facilitates its smoother passage across the bilayer. In contrast, ARC bears a carboxylic acid group that may be ionized under physiological conditions, increasing its polarity and reducing its compatibility with the hydrophobic membrane interior. This could contribute to the higher energy barrier and resistivity observed for ARC. Overall, these results suggest that subtle differences in polarity, ionization, and molecular structure can influence how these compounds interact with and traverse the BBB membrane.

**Table 2.** The Log P and BBB prediction of the tested small molecules.

| Compounds | log P | Predicted Log P<sub>eff</sub> | Experimental<br>BBB+/- | Predicted BBB+/- |
| --- | --- | --- | --- | --- |
| Antipyrine | -2.0 | -2.91 | Non-permeable | Non-permeable |
| Mannitol | -6.62 | -7.05 | Non permeable | Non-permeable |
| Ibuprofen | -0.63 | -0.88 | Permeable* | Non-Permeable |
| ARC | Data not available | -10.91 | Data not available | Non-permeable |
| CAPE | Data not available | 0.28 | Data not available | Permeable |

*Predicted BBB+ if $\log P_{eff} \geq 0$ , predicted BBB- if $\log P_{eff} < 0$ . \*Ibuprofen is the exception (Ooms et al. 2002)*

Although molecular modelling and simulations are the easier ways to study biological problems, and it is challenging to model existing systems, the critical components of the systems are chosen and simulated to get results similar to experimental data. In this study, we have not included all the complexities, like other lipids, proteins, and exporters in the BBB, to make the model simpler and easier to simulate. Likewise, the steered molecular dynamics simulation and the umbrella sampling have some limitations. They require the use of a restraint, or umbrella, to bias the system towards a specific region of the free energy landscape. This can introduce artefacts into the simulation, and the results may not be fully representative of the actual free energy of the system. Nevertheless, the studied model was well equilibrated, and the membrane properties were consistent with the previous data. The permeability values of the control molecules were also very close to the already published values. Therefore, the calculated BBB permeability of the natural compounds is expected to be close to their actual permeation.

## 4. Conclusions

The cell membrane permeability of a drug molecule is essential to interact with its target. In this study, we have modelled a BBB lipid bilayer by including all the essential lipid molecules known in the BBB to study the permeation of natural molecules, ARC and CAPE. The study showed that ARC had the high energy barrier hurdle to permeate through the BBB passively, while CAPE was mainly hindered by the polar head and mid-region of the lipids and had the very low resistance to pass through the BBB bilayer.

## Author’s Contribution

Conception and design- D.S. and VK.; Computational analysis- V.K.; Manuscript writing and Proofreading- V.K, S.C.K, R.W. and D.S; supervision- S.C.K, R.W. and D.S. All authors have read and agreed to the published version of the manuscript.

## References

Abraham, Mark James, Teemu Murtola, Roland Schulz, et al. 2015. “GROMACS: High Performance Molecular Simulations through Multi-Level Parallelism from Laptops to Supercomputers.” SoftwareX 1-2 (September): 19–25.

Allen, M. P., and D. J. Tildesley. 1987. Computer Simulation of Liquids. Clarendon Press.

Boggara, Mohan Babu, and Ramanan Krishnamoorti. 2010. “Partitioning of Nonsteroidal Antiinflammatory Drugs in Lipid Membranes: A Molecular Dynamics Simulation Study.” Biophysical Journal 98 (4): 586–595.

Campbell, Scott D., Karen J. Regina, and Evan D. Kharasch. 2014. “Significance of Lipid Composition in a Blood-Brain Barrier-Mimetic PAMPA Assay.” Journal of Biomolecular Screening 19 (3): 437–444.

Carpenter, Timothy S., Daniel A. Kirshner, Edmond Y. Lau, Sergio E. Wong, Jerome P. Nilmeier, and Felice C. Lightstone. 2014. “A Method to Predict Blood-Brain Barrier Permeability of Drug-like Compounds Using Molecular Dynamics Simulations.” Biophysical Journal 107 (3): 630–641.

Daneman, Richard, and Alexandre Prat. 2015. “The Blood-Brain Barrier.” Cold Spring Harbor Perspectives in Biology 7 (1): a020412.

Faramarzi, Sadegh, Marlene T. Kim, Donna A. Volpe, Kevin P. Cross, Suman Chakravarti, and Lidiya Stavitskaya. 2022. “Development of QSAR Models to Predict Blood-Brain Barrier Permeability.” Frontiers in Pharmacology 13 (October): 1040838.

Guixà-González, Ramon, Ismael Rodriguez-Espigares, Juan Manuel Ramírez-Anguita, et al. 2014. “MEMBPLUGIN: Studying Membrane Complexity in VMD.” Bioinformatics (Oxford, England) 30 (10): 1478–1480.

Hofsäss, Christofer, Erik Lindahl, and Olle Edholm. 2003. “Molecular Dynamics Simulations of Phospholipid Bilayers with Cholesterol.” Biophysical Journal 84 (4): 2192–2206.

Jo, Sunhwan, Taehoon Kim, Vidyashankara G. Iyer, and Wonpil Im. 2008. “CHARMM-GUI: A Web-Based Graphical User Interface for CHARMM.” Journal of Computational Chemistry 29 (11): 1859–1865.

Kaul, Ashish, Raviprasad Kuthethur, Yoshiyuki Ishida, Keiji Terao, Renu Wadhwa, and Sunil C. Kaul. 2021. “Molecular Insights into the Antistress Potentials of Brazilian Green Propolis Extract and Its Constituent Artepillin C.” Molecules (Basel, Switzerland) 27 (1): 80.

Kinnun, Jacob J., K. J. Mallikarjunaiah, Horia I. Petrache, and Michael F. Brown. 2015. “Elastic Deformation and Area per Lipid of Membranes: Atomistic View from Solid-State Deuterium NMR Spectroscopy.” Biochimica et Biophysica Acta 1848 (1 Pt B): 246–259.

Kulkarni, Namrata Pramod, Bhupesh Vaidya, Acharan S. Narula, and Shyam Sunder Sharma. 2021. “Neuroprotective Potential of Caffeic Acid Phenethyl Ester (CAPE) in CNS Disorders: Mechanistic and Therapeutic Insights.” Current Neuropharmacology 19 (9): 1401–1415.

Kumar, Shankar, John M. Rosenberg, Djamal Bouzida, Robert H. Swendsen, and Peter A. Kollman. 1992. “THE Weighted Histogram Analysis Method for Free-energy Calculations on Biomolecules. I. The Method.” Journal of Computational Chemistry 13 (8): 1011–1021.

Lee, Jumin, Xi Cheng, Jason M. Swails, et al. 2016. “CHARMM-GUI Input Generator for NAMD, GROMACS, AMBER, OpenMM, and CHARMM/OpenMM Simulations Using the CHARMM36 Additive Force Field.” Journal of Chemical Theory and Computation 12 (1): 405–413.

Luco, J. M. 1999. “Prediction of the Brain-Blood Distribution of a Large Set of Drugs from Structurally Derived Descriptors Using Partial Least-Squares (PLS) Modeling.” Journal of Chemical Information and Computer Sciences 39 (2): 396–404.

Marrink, Siewert-Jan, and Herman J. C. Berendsen. 1994. “Simulation of Water Transport through a Lipid Membrane.” The Journal of Physical Chemistry 98 (15): 4155–4168.

Mukhopadhyay, Parag, Hans J. Vogel, and D. Peter Tieleman. 2004. “Distribution of Pentachlorophenol in Phospholipid Bilayers: A Molecular Dynamics Study.” Biophysical Journal 86 (1 Pt 1): 337–345.

Ong, S., H. Liu, and C. Pidgeon. 1996. “Immobilized-Artificial-Membrane Chromatography: Measurements of Membrane Partition Coefficient and Predicting Drug Membrane Permeability.” Journal of Chromatography A 728 (1-2): 113–128.

Ooms, Frédéric, Peter Weber, Pierre Alain Carrupt, and Bernard Testa. 2002. “A Simple Model to Predict Blood-Brain Barrier Permeation from 3D Molecular Fields.” Biochimica et Biophysica Acta 1587 (2-3): 118–125.

Orsi, Mario, Wendy E. Sanderson, and Jonathan W. Essex. 2009. “Permeability of Small Molecules through a Lipid Bilayer: A Multiscale Simulation Study.” The Journal of Physical Chemistry. B 113 (35): 12019–12029.

Ottaviani, Giorgio, Sophie Martel, and Pierre-Alain Carrupt. 2006. “Parallel Artificial Membrane Permeability Assay: A New Membrane for the Fast Prediction of Passive Human Skin Permeability.” Journal of Medicinal Chemistry 49 (13): 3948–3954.

Park, Sanghyun, and Klaus Schulten. 2004. “Calculating Potentials of Mean Force from Steered Molecular Dynamics Simulations.” The Journal of Chemical Physics 120 (13): 5946–5961.

Petrache, H. I., S. W. Dodd, and M. F. Brown. 2000. “Area per Lipid and Acyl Length Distributions in Fluid Phosphatidylcholines Determined by (2)H NMR Spectroscopy.” Biophysical Journal 79 (6): 3172–3192.

Radhakrishnan, Navaneethan, Sunil C. Kaul, Renu Wadhwa, and Durai Sundar. 2022. “Phosphatidylserine Exposed Lipid Bilayer Models for Understanding Cancer Cell Selectivity of Natural Compounds: A Molecular Dynamics Simulation Study.” Membranes 12 (1): 64.

Shahane, Ganesh, Wei Ding, Michail Palaiokostas, and Mario Orsi. 2019. “Physical Properties of Model Biological Lipid Bilayers: Insights from All-Atom Molecular Dynamics Simulations.” Journal of Molecular Modeling 25 (3): 76.

Sharifian Gh Mohammad. 2021. “Recent Experimental Developments in Studying Passive Membrane Transport of Drug Molecules.” Molecular Pharmaceutics 18 (6): 2122–2141.

Smith, David J., Jeffery B. Klauda, and Alexander J. Sodt. 2019. “Simulation Best Practices for Lipid Membranes [article v1.0].” Living Journal of Computational Molecular Science 1 (1): 5966.

Stamatovic, Svetlana M., Richard F. Keep, and Anuska V. Andjelkovic. 2008. “Brain Endothelial Cell-Cell Junctions: How to ‘Open’ the Blood Brain Barrier.” Current Neuropharmacology 6 (3): 179–192.

Sun, Duxin, Wei Gao, Hongxiang Hu, and Simon Zhou. 2022. “Why 90% of Clinical Drug Development Fails and How to Improve It?” Acta Pharmaceutica Sinica. B 12 (7): 3049–3062.

Tewes, B. J., and H. J. Galla. 2001. “Lipid Polarity in Brain Capillary Endothelial Cells.” Endothelium: Journal of Endothelial Cell Research 8 (3): 207–220.

Toropov, Andrey A., Alla P. Toropova, Marten Beeg, Marco Gobbi, and Mario Salmona. 2017. “QSAR Model for Blood-Brain Barrier Permeation.” Journal of Pharmacological and Toxicological Methods 88 (Pt 1): 7–18.

Torrie, G. M., and J. P. Valleau. 1977. “Nonphysical Sampling Distributions in Monte Carlo Free-Energy Estimation: Umbrella Sampling.” Journal of Computational Physics 23 (2): 187–199.

Vanommeslaeghe, K., E. Hatcher, C. Acharya, et al. 2010. “CHARMM General Force Field: A Force Field for Drug-like Molecules Compatible with the CHARMM All-Atom Additive Biological Force Fields.” Journal of Computational Chemistry 31 (4): 671–690.

Wadhwa, Renu, Neetu Singh Yadav, Shashank P. Katiyar, et al. 2021. “Molecular Dynamics Simulations and Experimental Studies Reveal Differential Permeability of Withaferin-A and Withanone across the Model Cell Membrane.” Scientific Reports 11 (1): 2352.

Zhang, Liying, Hao Zhu, Tudor I. Oprea, Alexander Golbraikh, and Alexander Tropsha. 2008. “QSAR Modeling of the Blood-Brain Barrier Permeability for Diverse Organic Compounds.” Pharmaceutical Research 25 (8): 1902–1914.

